# Social Perception and Interaction Database – a novel tool to study social cognitive processes with point-light displays

**DOI:** 10.1101/729996

**Authors:** Ł. Okruszek, M. Chrustowicz

**Affiliations:** Social Neuroscience Lab, Institute of Psychology, Polish Academy of Sciences; Faculty of Psychology, University of Warsaw

## Abstract

**Introduction:** The ability to detect and interpret third-party encounters (TPE) is one of the crucial skills enabling people to operate in the social world. Multiple lines of evidence converge towards the preferential processing of TPE when compared to the individual actions of multiple agents, even if the actions of agents were visually degraded to minimalistic point-light displays (PLDs). Here, we present a novel PLD dataset (Social Perception and Interaction Database; SoPID) that may be used for studying multiple levels of social information processing.

**Methods:** During a motion-capture session, two pairs of actors were asked to perform a wide range of dyadic 3-second actions, including: (1) neutral, gesture-based communicative interactions (COM); (2) emotional exchanges (Happy/Angry); (3) synchronous physical activity of actors (SYNC); and (4) independent actions of agents, either object-related (ORA) or non-object related (NORA). The stimuli were then transformed into PLDs. Two validation studies (each with 20 healthy individuals) were then performed to establish the recognizability of the SoPID vignettes.

**Results:** The first study showed a ceiling level accuracy for discrimination of communicative vs. individual actions (93% +/- 5%) and high accuracy for interpreting specific types of actions (85 +/- 4%) from the SoPID. In the second study, a robust effect of scrambling on the recognizability of SoPID vignettes was observed in an independent sample of healthy individuals.

**Discussion:** These results suggest that the SoPID may be effectively used to examine processes associated with communicative interactions and intentions processing. The database can be accessed via Open Science Framework (https://osf.io/dcht8/).

## Introduction

Multiple lines of evidence converge towards the preferential processing of encounters between other agents. Communicative interactions have been shown to be easily discriminated from other types of dyadic actions (Manera, Becchio, Schouten, Bara, & Verfaillie, 2011; Manera et al., 2015; Manera, Schouten, Becchio, Bara, & Verfaillie, 2010), gain preferential access to awareness (Su, van Boxtel, & Lu, 2016), and are encoded as a single unit in working memory (Ding, Gao, & Shen, 2017). Psychophysics experiments have also shown that healthy individuals are able to utilize top-down knowledge about the communicative gesture of one agent to predict both the type (Manera, Del Giudice, Bara, Verfaillie, & Becchio, 2011) and timing (Manera, Schouten, Verfaillie, & Becchio, 2013) of the other agent’s response. Furthermore, social interactions processing elicits widespread activation of the main “social brain” networks, compared to the individual actions of multiple agents (Centelles, Assaiante, Nazarian, Anton, & Schmitz, 2011; Isik, Koldewyn, Beeler, & Kanwisher, 2017; Quadflieg & Koldewyn, 2017; Walbrin, Downing, & Koldewyn, 2018). Importantly, these effects may be observed for both naturalistic full displays of agents (Canessa et al., 2012; Quadflieg, Gentile, & Rossion, 2015; Van den Stock, Hortensius, Sinke, Goebel, & de Gelder, 2015; Wang, Huang, Zhang, Zhang, & Cacioppo, 2015) and minimalistic point-light displays of social interactions (Centelles et al., 2011; Isik et al., 2017; Petrini, Piwek, Crabbe, Pollick, & Garrod, 2014; Walbrin et al., 2018). Developed by Johansson (Johansson, 1973), point-light methodology limits the presentation of the agents to a set of light-dots representing the head, limbs, and major joints of the agent’s body. Despite the extremely limited amount of visual information presented via point-light displays (PLDs), this type of vignette has been shown to carry enough information to enable the recognition of an agent’s action (Dekeyser, Verfaillie, & Vanrie, 2002; Vanrie & Verfaillie, 2004), affective state (Vaskinn et al., 2015), and a wide range of physical characteristics (Troje et al., 2013). Furthermore, point-light stimuli have also been used to investigate communicative intentions processing from both single (Zaini, Fawcett, White, & Newman, 2013) and dyadic displays (Manera et al., 2010).

Manera et al. (Manera et al., 2010) presented the Communicative Interaction Database (CID) – a set of 20 stimuli that presents dyadic interactions based on the stereotypical use of communicative gestures with point-light motion. CID stimuli have been used to examine both reflective (Manera et al., 2015) and reflexive (Manera, Becchio, et al., 2011) social cognitive processes in healthy individuals. Stimuli from the CID have also been used to create a multilingual task for studying communicative interaction recognition (Manera et al., 2015), which has been effectively applied to study social cognition across various clinical populations (patients with schizophrenia (Okruszek et al., 2015; Okruszek, Piejka, Wysokiński, Szczepocka, & Manera, 2018), high functioning individuals with autism spectrum disorders (von der Lühe et al., 2016), patients with temporal lobe epilepsy (Bala et al., 2018)). Furthermore, CID stimuli have been applied to investigate the neural correlates of communicative processing (Isik et al., 2017; Walbrin et al., 2018).

At the same time, given the widespread nature of third party encounters (TPE) processing across neural networks, a recent review of neural and behavioral findings in this area concluded that the development of TPE localizers, which entail various types of social interaction vignettes, may facilitate research in this area (Quadflieg & Koldewyn, 2017). Studies based on static pictures of various types of social interactions have observed differential patterns of brain activity (Canessa et al., 2012) and connectivity (Arioli et al., 2017) in affective vs. cooperative interactions. Yet, due to the limited availability of point-light stimuli, previous studies on TPE processing from PLDs either pooled various types of communicative interactions into one category (e.g. (Centelles et al., 2011)) or presented only certain types of interactions (usually encounters based on the typical use of communicative gestures: (Isik et al., 2017; Walbrin et al., 2018)). Furthermore, it has been shown that communicative intentions may be differentially processed from the second person (receiver) and third person (observer) perspective (Ciaramidaro, Becchio, Colle, Bara, & Walter, 2014). Thus, to address *second person neuroscience* postulates (Schilbach et al., 2013), future studies should compare the processing of communicative intentions both from second person (single walker presenting gesture towards observer) and third person (dyadic displays of agents acting towards each other) perspectives. The aim of the current project was to develop a database of point-light stimuli (Social Perception and Interaction Database; SoPID) that addresses the abovelisted issues by allowing one to create point-light vignettes with a wide range of communicative and individual actions while flexibly manipulating the number of agents presented (one vs two), the viewing perspective, and display options.

## Methods

### Pre-capturing session

Two pairs of professional actors took part in the motion capture procedure. One dyad consisted of male actors and one of female actresses. During the pre-capturing session, actors were familiarized with the list of actions that were to be recorded. The list of situations to be recorded consisted of six categories, each with 5-10 situations (see Appendix A for a full list of the SoPID stimuli). For the communicative interactions (COM), each of the actors was asked to play both the person initiating the interaction via a communicative gesture (Agent A) and the person responding to the communicative gesture of the other agent (Agent B), thus producing two different takes on each COM situation. A short description of each action was provided to ensure that each dyad was enacting similar communicative intention and a similar behavioral response. To ensure the temporal synchronization of the animations, three sound signals were presented during each recording: first to signal the onset of the recording (played at T = 0 s.), second to signal the half-time of the recording (played at T = 1.5 s.), and third to signal the end of the recording (played at T = 3 s.). For animations that presented the sequential actions of both agents (e.g. Agent A asks Agent B to stand up, Agent B stands up), the actor playing Agent A was asked to start his action at T = 0, while the actor playing Agent B was asked to start responding at T = 1.5 s. The situations were rehearsed until both actors were able to perform them with the required timing. Moreover, to ensure the naturalistic yet expressive movement of the actors during the capturing sessions, a professional choreographer oversaw the actors’ rehearsal during the pre-capturing session.

### Motion capture

The motion capture session was performed via a motion-capture studio (*White Kanga* studio; Warsaw) using an OptiTrack (NaturalPoint, Corvallis, OR, USA) motion tracking system. Twelve OptiTrack Prime 13 cameras were utilized to record the movements of the actors at a 120 Hz rate. The actors wore 41 reflective spherical markers placed according to the OptiTrack Baseline+Hinged Toe system (full list and anatomical locations of the markers available at https://v20.wiki.optitrack.com/index.php?title=Baseline_%2B_Hinged_Toe,_with_Headband_(41)). The motion capture room was a 7 × 7 meters square with a 3.8 meter high; a white line was painted on the floor of the motion capture room to mark each actor’s subspace (7 × 3.5 meter) within the 7 × 7 meter motion capture box. With the exception of animations that included physical contact between the agents, the actors were asked to confine their actions within their subspaces. Similarly, most of the sequences were recorded with actors facing each other at a proximity of around 3 meters. At the beginning and end of each recording, the actors were asked to perform a T-pose (reference pose) at the central position of their subspace. Additional props were used for the sequences that included object-related actions (i.e. shovel, carton box, axe, saw, broom, glass, hammer, toothbrush, football, chair). No markers were used to tag the prop positions during the session. For the animations that presented interaction between the agents (communicative interactions, happy/angry and synchronous activity), the actions of both agents were recorded simultaneously to ensure that the response of one agent was congruent with the action of the other agent in terms of position, proxemics, and timing. Actions for object- and non-object related displays were recorded individually to minimize the potential effects of between-agents synchrony while performing the actions. Similarly, as during the pre-capturing session, sound cues were used to inform actors about beginning (T = 0 s.), middle (T 1.5 s), and end (T = 3 s.) points of each three-second period.

**Figure 1.**
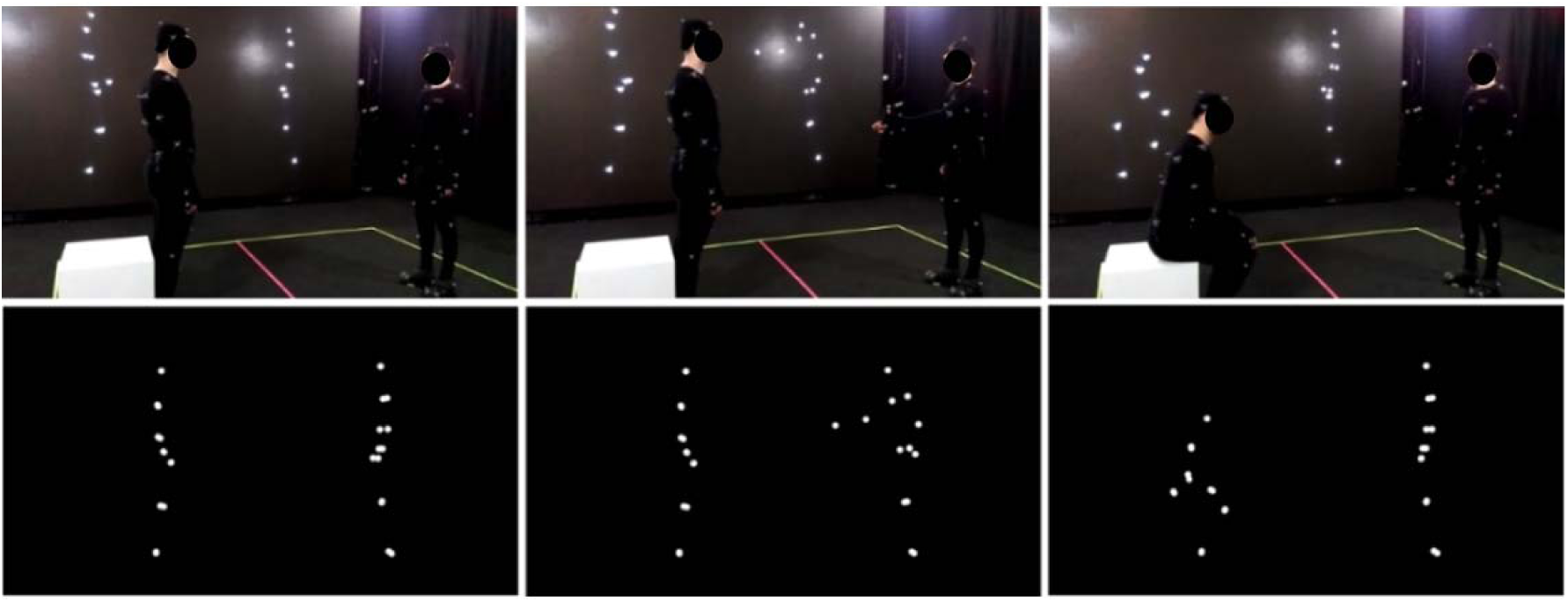
Original (upper) and PLD (lower) version of the item presenting communicative interaction from SoPID (A (on the right) asks B (on the left) to sit down; B sits down)

### Data processing

Data from the motion capture session were further processed using OptiTrack Motive 1.9 beta software. 2-D data from twelve cameras were used to obtain the 3-D coordinates of each marker. Skeleton models consisting of 13 bright dots corresponding to the head, arms, elbows, wrists, hips, knees, and ankles of each actor were animated. Data preprocessing included inspection of each of the recordings, data trimming to the period between the onset (T = 0 s.) and offset (T = 3 s.) of the action, and manual smoothing in case of any vibrating or fluttering movements. The preprocessed data were extracted to FBX files.

### Social Perception and Interaction Database

To enable users without programming skills to access and customize the stimuli according to their needs, preprocessed stimuli may be accessed via an interface that is based on the Unity engine (SoPID). The SoPID interface allows for modification of numerous stimuli characteristics and exports the customized stimuli to movie files (.mp4) using the FFmpeg codec. Overall, 64 different actions of each agent can be accessed via the SoPID and used to create experimental stimuli. Each of the recorded actions may be accessed either separately as a solo action or merged with a second action to produce a vignette presenting a pair of agents. This way, the SoPID allows one to produce a wide range of vignettes presenting a single agent’s communicative or individual actions, as well as to combine the actions of two agents to obtain either congruent (by selecting one out of four Agent A’s ‘Communicative Gestures’ and any of the corresponding responses of Agent B or by using a combination of either ‘Happy’, ‘Angry’ or ‘Synchronous Activity’ actions) or incongruent (e.g. by mixing Agent A’s communicative action with a non-corresponding action of Agent B) social interactions or dyadic individual actions of agents.

The SoPID interface also allows one to flexibly adjust camera position. Four standard camera positions may be selected, with the ‘Front’ position corresponding to a 270 degree display from the CID (Agent A on the left and Agent B on the right) with the camera being place on the middle line between the agents, at the one meter height and 15 meters from the agents. Furthermore, by using the ‘Free’ option, one may fully customize both the camera placement (x – left/right; y – up/down; z – closer/further; values in meters) and rotation (x – up/down; y – left/right; z – horizontal/vertical; values in degrees). Default settings for the four default camera positions are presented in Supplementary Table 1. Both ortographic and perspective projections may be used to manipulate the availability of depth cues in the animations. Additionally, marker size may be changed (‘Marker size’, values in centimeters) to modify the agents’ appearances and stick figures can be created instead of point-light displays (‘Show skeleton’). Finally, two standard modifications that are commonly used in point-light studies can be applied directly via the SoPID. First, by using the ‘Flicker’ option, one may selectively limit the visual availability of the stimuli by selecting the maximal number of simultaneously displayed markers (0-13) and the time range for marker display/ disappearance. Markers are flickered by randomly assigning the onset and offset time values separately for each marker with regard to the time range provided by the user. In addition, by using the ‘Scramble’ option one may spatially scramble the initial spatial position of each marker. Scrambling is applied by randomly drawing one out three dimensions for each marker and relocating its initial position by X centimeters from its initial position in the selected direction (e.g. 100 % scrambling moves each marker by one meter in either the x, y or z dimension). ‘Flicker’ and ‘Scramble’ can be applied to both agents or selectively to each agent. The database can be accessed via Open Science Framework (https://osf.io/dcht8/).

**Figure 2.**
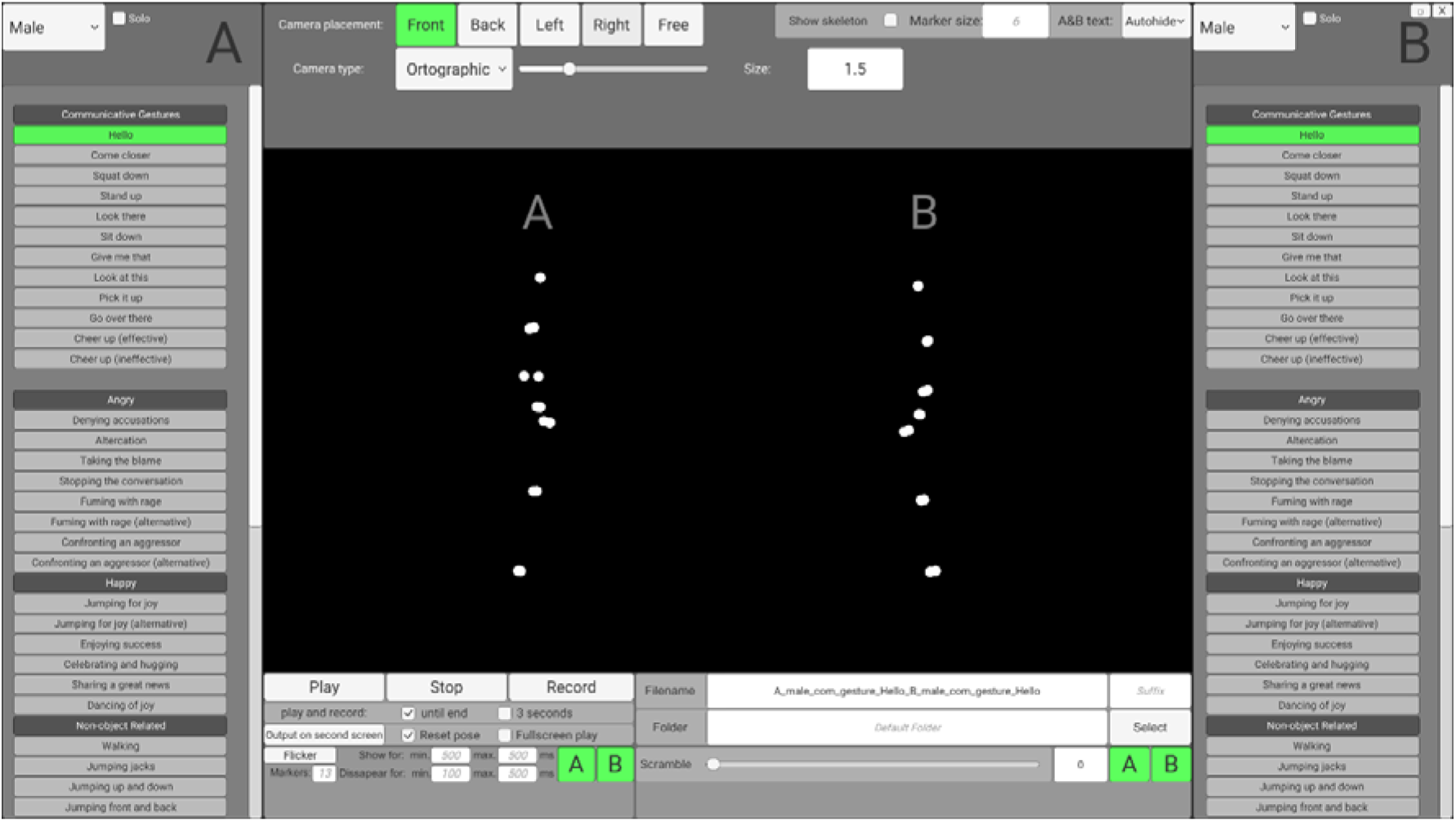
Social Perception and Interaction Database interface.

### Validation studies

To examine the recognizability of the presented actions and the effectiveness of the scrambling mechanism, two SoPID validation studies were performed. Participants for each of the studies were recruited from the students of Warsaw-based universities. Participants were tested individually, and had not participated in point-light experiments prior to the examination.

### Study 1

Twenty right-handed participants (9M/11F; 25.9 +/- 9.1 yrs. old) completed Study 1. Stimuli: Fifty-seven animations presenting the actions of two agents were created using the SoPID (perspective camera with FoV = 10°, camera position = front, and marker size = 6). Six types of animations were presented throughout the study: Communicative gestures [COM, 10 animations: “Hello” (Female 2 as Agent A); “Come closer” (Male 1 as Agent A), “Squat down” (F2), “Stand up” (M1), “Look there” (F1), “Sit down” (M2), “Give me that” (F2), “Look at this” (M1), “Pick it up” (F2), “Go over there” (M2)]; Angry exchanges (Angry, 5 animations: “Denying accusations”, “Taking the blame”, “Stopping the conversation”, “Fuming with rage”, “Confronting an aggressor (alternative)”); Happy exchanges (Happy: 5 animations: “Jumping for joy”, “Enjoying success”, “Celebrating and hugging”, “Sharing a great news”, “Dancing of joy”); Non-object related dyadic individual actions (NORA, 10 animations: “Walking”, “Jumping jacks”, “Jumping up and down”, “Jumping front and back”, “Arm waving”, “Hip swinging”, “Torso twist”, “A-Skip”, “Squat down”, “Lateral step”, “Lateral kick”); Object related dyadic individual actions (ORA, 9 animations: “Shoveling”, “Lifting the box”, “Chopping wood”, “Sawing”, “Digging”, “Sweeping the floor”, “Drinking”, “Hammering a nail”, “Brushing teeth”) and Synchronous activity of two agents [SYNC, 8 animations: “Dancing” (M/F), “Fencing” (M), “Football” (F), “Throwing the ball” (M), “Boxing” (F), “Kickboxing” (M/F)]. To increase the comparability of recognition accuracy levels across the categories, stimuli from the Angry and Happy categories were duplicated by presenting each situation twice, with either male or female actors. ORA and NORA movies were created by merging the displays of two different actions performed by two same-sex actors. Displays of each set of actions with either male or female actors were included, thus producing 11 NORA and 9 ORA movies in total.

#### Task

Each stimulus was presented twice, after which participants were asked to: (1) classify the presented action as an interaction (behavior of one agent affects the behavior of the other) or the individual actions of agents, and (2) to provide a verbal description of the actions of the agents. The order of the stimuli presentation was pseudorandomized to avoid subsequent presentation of more than two stimuli from the same category. The paradigm was programmed using NBS Presentation 20, and the whole procedure took approximately one hour. Verbal descriptions provided by the participants were scored by a rater who did not participate in data collection. Spontaneous descriptions for COM, SYNC, Happy, and Angry were scored in a dichotomic manner (2 points for a correct verbal description vs 0 points for an incorrect description). Accuracy for ORA and NORA stimuli was calculated by scoring one point for each correctly recognized action from male and female presentations (0-2 points). For interaction vs. individual actions classification, COM, Angry, Happy and SYNC were treated as falling into the category ‘interaction’, while ORA and NORA were treated as ‘individual actions.’ Two items (‘Dancing for joy’ and ‘Fuming with rage’) without any explicit communicative cues were discarded from this part of the analysis. To examine between-category differences in accuracy levels, one way ANOVAs with Type of animation (six levels) were performed separately for interaction recognition and spontaneous identification of actions.

#### Results

##### Recognition of communicative intentions

No between category differences were observed for classifying actions as either communicative or individual (F(5, 15) = 1.3; n.s., η_p_^2^=0.07), with ceiling level recognition for all types of items [Angry: 95% +/- 9%; Happy: 89% +/- 13%; COM: 95% +/- 10%; SYNC: 91% +/- 11%; NORA: 94% +/- 10%; ORA: 93% +/- 8%].

##### Identification of specific action

A main effect of category was observed for the accuracy of identification of specific actions (F(5,15) = 23.9, p < 0.001, η_p_^2^=0.56). Further investigation of this effect revealed the highest recognition for SYNC (96% +/- 6%), NORA (95% +/- 5%) and Happy (92% +/- 11%), each of which were identified at higher level than Angry (81% +/- 12%) and COM (78% +/- 12%). Furthermore, actions from all of the categories were identified more accurately than ORA (67% +/- 13%).

### Study 2

Twenty right-handed participants (10M/10F; 24.2 +/- 7.7 years old) completed Study 2. Stimuli: Twenty animations [‘come closer’ (F), ‘squat down’ (M), ‘stand up’ (M, F), ‘go over there’ (F), ‘altercation’ (M, F), ‘jumping for joy’ (F), ‘denying accusations’ (M), ‘jumping for joy (alternative)’ (M), ‘walking’ (F), ‘lateral kick’ (F), ‘hip swinging’ (M), ‘A-skip’ (M), ‘squat down’ (M), ‘lifting the box’ (F), ‘sweeping the floor’ (F), ‘brushing teeth’ (F), ‘chopping wood’ (M), ‘digging’ (M)] presenting the action of a single agent (Agent A in case of COM, Angry and Happy) were created from the SoPID (ortographic camera (size = 1.5), camera position = right, and marker size = 6). Each animation was rendered at four scrambling levels: 0%, 15%, 30%, and 100%. Thus, 80 animations were presented during the experimental procedure.

#### Task

Upon presentation of each animation, participants were asked to indicate whether the presented animation resembled human motion. Completion of the whole experimental procedure took approximately 20 minutes. The number of stimuli classified as ‘human’ at each scrambling level was compared to examine the effectiveness of the scrambling procedure. The results were analyzed using rmANOVA with the within-subject factor Scrambling (4 levels: 0%, 15%, 30%, 100%).

#### Results

Due to the strongly outlying results, one participant was excluded from the analysis. A robust effect of scrambling was observed (F(3, 16) = 400.9; p < 0.001, η_p_^2^=0.96). Unscrambled stimuli were classified significantly more often as human motion (98% +/- 3%) when compared to 15% (68% +/- 18%), 30%- (20% +/- 13%) and 100% scrambled (3% +/- 5%) motion. Similarly, 15% stimuli were classified as human motion more often than 30% and 100% scrambled displays, and 30% scrambled stimuli were classified as human more often than 100% scrambled displays. All of the contrasts were significant at p < 0.001.

## Discussion

The present paper describes the Social Perception and Interaction Database, a novel set of point-light displays that enables study of the processing of a wide range of communicative and individual actions from single-agent and dyadic vignettes. The SoPID includes 32 vignettes presenting various types of social interactions between two agents, including standard use of communicative gestures (COM), synchronous physical activity (SYNC) and affective exchanges (either Happy or Angry), as well as 20 animations of each actor performing either object- (ORA) or non-object-related (NORA) individual actions. Moreover, by allowing users to create vignettes by mixing the actions of both agents, both within each category and between the categories, a wide range of novel stimuli can be created to study the processing of typical and atypical social interactions.

Previous studies that used the CID database showed high accuracy in recognition of communicative vs. individual dyadic actions in healthy individuals (Manera et al., 2015; Manera, von der Lühe, Schilbach, Verfaillie, & Becchio, 2016). Similarly, we observed ceiling level accuracy for classifying stimuli as either communicative or individual across the six categories of stimuli included in the first of the validation studies (ranging from 89% for Happy to 95% for COM). Furthermore, the accuracy of identification of specific communicative actions from the COM category (78% +/- 12%) was at a similar level as reported previously for the multilingual CID task (74% +/- 18%; Manera et al., 2016).

Interestingly, more accurate identification of specific actions was observed for three other categories of stimuli included in the study (Happy, NORA, SYNC). This result may be linked to the fact that both NORA and SYNC vignettes presented low-level physical activity that is usually associated with whole-body motion (e.g. jumping or kick-boxing). Thus, (1) they did not require intention attribution and therefore could have been described in a less complex form, and (2) presented more whole-body cues compared to COM, which usually presented the sequential actions of agents initiated by a hand motion representing the communicative gesture of Agent A.

Furthermore, while no accuracy differences were observed between the recognition of neutral communicative interactions (COM) and negatively-valenced social exchanges (Angry), positively-valenced social exchanges (Happy) were identified more effectively than COM and Angry vignettes. This finding is in line with Lee and Kim (Lee & Kim, 2016), who recently showed that the positive emotional valence (happiness) of stimuli may facilitate biological motion processing. Furthermore, Piwek et al. (Piwek, Pollick, & Petrini, 2015) also found greater recognition of happy compared to angry dyadic interactions presented with point-light displays, especially at the low and moderate levels of emotion intensity.

Finally, we observed that object-related individual actions were identified less accurately than any other type of SoPID animations. While ORA were classified as an individual actions with high accuracy (93% +/- 8% classification rate), identification of specific object-related actions was observed to be more challenging than for any other category of stimuli from the SoPID (67% +/- 13%). A recent study with 24 healthy individuals found slightly higher accuracy for the description of object-related individual actions presented with single-agent point-light displays (74% +/- 11%; (Jaywant et al., 2016)); however, the interpretation of the actions in the study may have been facilitated by the use of point-light displays with high-resolution representations of the hands and fingers (additional 10 point-lights; (Zaini et al., 2013)). Still, the spontaneous un-cued recognition of ORA stimuli from standard 13-dot PLDs created with the SoPID was at a similar level as the recognition rates of individual actions from the well-established stimuli set by Vanrie et al. (Vanrie & Verfaillie, 2004), which reported a mean accuracy rate of 63% for a set of individual point-light actions in 11 observers producing spontaneous descriptions of the vignettes.

Our second study aimed to test an additional feature of the SoPID, i.e. a scrambling mechanism. We observed robust scrambling effects: unscrambled and 100% scrambled stimuli were almost unanimously categorized as, respectively, human and non-human motion. Furthermore, more subtle effects of scrambling were also observed: a significant portion of the 15% scrambled stimuli were classified by participants as resembling human motion, while a large majority (80%) of the 30% scrambled stimuli were classified as non-human motion.

Taken together, these results suggest that SoPID stimuli may be effectively used in a wide range of experiments examining both the basic (e.g. recognition of biological vs. scrambled motion) and higher-order (e.g. recognition of communicative intentions of affective states from PLDs) processing of biological motion. By providing multiple ways to customize the PLD vignettes, the SoPID may be effectively used to examine a wide range of questions regarding the social cognitive mechanisms associated with communicative interactions processing. For example, by presenting the same stimuli from a second- and third-person perspective, one may examine the impact of the participant vs. observer role for communicative intentions processing. Furthermore, by manipulating the congruency of the actions in dyadic displays, the SoPID allows one to address the issue of the role of the second agent’s response for processing of communicative interactions. Finally, the necessity of developing new localizers for studying the neural correlates of third party encounter processing has been recently stressed (Quadflieg & Koldewyn, 2017). By providing multiple different options for stimuli presentation (e.g. point-light agents vs. stick figures) and a wide range of available actions from various semantic categories, the SoPID allows one to create enough stimuli to develop such localizers for neuroimaging and neurophysiological research.

## Acknowledgements

This work was supported by the National Science Centre, Poland (Grant No: 2016/23/D/HS6/02947). We would like to thank to the Bartłomiej Leszek, Marcin Kocik and White Kanga studio personnel, as well as Aleksandra Matlingiewicz, Sonia Jachymiak, Piotr Sedkowski, Piotr Watroba and Artur Marchlewski and Grzegorz Pochwatko for their help in stimuli creation.

